# Elevated temperature enhances the infectivity of a pathogenic noncoding RNA and the likelihood of its host expansion

**DOI:** 10.1101/2025.09.19.677415

**Authors:** Jie Hao, Zongyun Yan, Yunhan Wang, Junfei Ma, Benjamin A. Merritt, Ethan Tucker, Amit Levy, Bin Liu, Svetlana Y. Folimonova, Jeongim Kim, Zhonglin Mou, Yiliang Ding, Ying Wang

**Author notes:** Correspondence (J.H.), (J.M.), (Y.D.), (YingW.). These authors contributed equally (Jie Hao, Zongyun Yan, Yunhan Wang, Junfei Ma).

## Abstract

Plant pathogens threaten global food security and cause substantial annual economic losses. Of particular concern are host jumps, in which pathogens expand their host range to new species, often resulting in sudden and difficult-to-control disease outbreaks. However, the ability to anticipate such events remains limited by an incomplete understanding of pathogen adaptability and the mechanisms governing nonhost resistance. Here, we show that temperature modulates the infectivity and host range of potato spindle tuber viroid (PSTVd), a pathogenic noncoding RNA that severely impacts solanaceous crops such as potato and tomato. Using *in vivo* single-molecule RNA structure probing, we found that higher temperatures drive PSTVd into a more uniform RNA conformation associated with enhanced infectivity in tomato. Concurrently, elevated temperature suppresses key host defense pathways, including RNA silencing and NPR1-mediated immunity. Notably, these conditions enable PSTVd to overcome nonhost resistance and establish infection in *Arabidopsis thaliana*. Our findings reveal that elevated temperature promotes PSTVd host expansion through multiple coordinated mechanisms, linking changes in pathogen behavior with increased host susceptibility, and suggest that ongoing climate warming may heighten the risk of viroid host jumps.

## Introduction

Plants are constantly challenged by diverse pathogens, including viruses and viroids. During pathogen attack, plants deploy multiple defense mechanisms, including innate immunity and RNA silencing (*1, 2*). Notably, pathogens often spill over to nonhost plants, but these spillover events mostly result in dead-end infection that causes no damage because of nonhost resistance (*3*). Nonhost resistance is a phenomenological rather than mechanistic concept, and the underlying mechanisms are often unique to specific pathogen-plant species combinations (*4*). Occasionally, new pathogen variants can emerge during spillovers, and nonhost plants could then become new hosts (*3*). Such host jumps can lead to outbreaks of new plant diseases (*5*). Ongoing climate change has been associated with increased outbreaks of viral and viroid diseases in agriculture (*6–8*). Climate change has been considered to influence growth conditions of crop hosts, the abundance and distribution of viruses and viroids in wild reservoir and/or vectors, and the rate of the viral pathogen spread (*8*). However, how environmental temperature modulates viral and viroid infectivity and host range remains poorly understood.

Viroids, single-stranded circular noncoding subviral RNAs, mainly infect crops and often cause significant yield losses (*9, 10*). Due to the noncoding nature, viroids must exploit host factors to accomplish the replication cycle and evade host defenses (*11*). For instance, potato spindle tuber viroid (PSTVd; *Pospiviroid fusituberis*; the type species of the family *Pospiviroidae*) utilizes its C-loop RNA motif to recruit host protein Virp1 and form the RNA-protein complex (RNP). This RNP then uses an importin alpha4 (IMPA4)-based nuclear import pathway for nuclear entry (*12*). In the nucleus, PSTVd forms another RNP with a splicing variant of transcription factor IIIA (TFIIIA-7ZF) to recruit RNA polymerase II (Pol II) for transcription (*13, 14*). This Pol II complex is remodeled from a 12-subunit into a 7-subunit complex adaptated for RNA-templated transcription (*15, 16*). The RNA intermediates produced by the remodeled Pol II are cleaved by an RNase III-type enzyme (*17*) and circularized by DNA ligase I (*18*). To optimize TFIIIA-7ZF level for replication, PSTVd modulates splicing by directly interacting with RPL5 (*19*), a splicing regulator of TFIIIA (*20*). All of the described host factors (Virp1, IMPA4, Pol II, TFIIIA-7ZF, DNA ligase I, RPL5) are highly conserved in the land plants (*21*), at least in *Arabidopsis thaliana*, which is a well-known nonhost plant that does not support systemic infection of any viroid (*22*). Yet, multiple viroids can still replicate in *A. thaliana* when expressed transgenically or introduced into protoplasts by transfection (*22–24*), suggesting that this nonhost nature is unlikely due to the absence of host factors.

The molecular basis underlying plant defense against viroids, particularly members in the *Pospiviroidae* family that replicate in the nucleus, is not fully understood. The RNA silencing machinery was regarded as the primary defense against viroids (*9*), mainly relying on Dicer-like 2 (DCL2) (*25, 26*). Nevertheless, transcriptomic analyses of multiple viroid-host pathosystems have revealed the activation of immune-related marker genes, such as Pathogenesis-Related 1. (*9, 27*). Whether plant innate immunity restricts viroid infection remains elusive. Thus, additional investigations are needed to further dissect the plant defense mechanisms in plant-viroid interactions.

Interestingly, reports of viroid jumps into new hosts have increased in the 21^st^ century (*28*). For instance, PSTVd was recently found in various solanaceous hosts other than potato and tomato (*29*). Moreover, the PSTVd host jump to dahlia, a non-solanaceous host, was documented during the past decade (*29*). Viroid jumps to new hosts may not always lead to severe plant diseases, but they nevertheless increase risks to agricultural production. For instance, the host jump of hop latent viroid to cannabis in recent years has caused severe symptoms and production losses in this new host (*30*). To date, the underlying mechanisms of viroid host range expansion remain unclear, hindering the ability to forecast and prepare for new viroid disease outbreaks.

Here, we show that PSTVd accumulated to higher titers and triggered more pronounced symptoms in tomato plants growing at 28°C, in comparison to the infection at 22°C. By comparing PSTVd titers in tomato mutant lines with loss-of-function of RNA silencing components, the data showed that the RNA silencing components exhibited distinct activities on PSTVd infection at different temperatures. Therefore, this temperature-dependent infectivity offers an opportunity to dissect new mechanisms restraining PSTVd infection. *In vivo* single-molecule RNA structurome (smStructure-Seq) analysis demonstrated that PSTVd RNAs fold into a more uniform structure at 28°C, in favor of entry into the replication site, suggesting the interaction between thermodynamic folding and environmental temperature facilitating viroid infection. Furthermore, NONEXPRESSOR OF PATHOGENESIS-RELATED GENES 1 (NPR1) was identified as a novel antiviroidal factor, and its activity against PSTVd was impaired at 28°C in tomato. The impaired NPR1 activity was linked to its failure in nuclear accumulation, although the salicylic acid level was induced in infected plants. Notably, exogenously applying a SA analog (acibenzolar-S-methyl) still achieved a high successful rate in blocking PSTVd infection in inoculated tomato plants by triggering the degradation of the required host factor and enzyme for PSTVd replication, suggesting the presence of multiple pathways in SA-mediated antiviroidal defense. Interestingly, high temperature also permitted PSTVd infection in a nonhost plant *A. thaliana*, and its titer was further increased in the *npr1* loss-of-function plants. Altogether, our results uncovered multi-layer regulations that promote PSTVd infectivity at 28°C, with the thermodynamic RNA folding and NPR1 activity as novel regulation in plant-viroid interactions. This study highlights the pressing risk of viroid host jumps during global climate changes, but also pinpoints the direction for developing sustainable measure to combat PSTVd infection.

## Results

### PSTVd infectivity is modulated by temperature

It has been noticed that higher temperature has an impact on the infection of plant viruses and viroids (*31–33*). In general, viral pathogens replicate to higher titers at higher temperatures. This effect is currently attributed to the activity of RNA silencing (*31–33*). In our experimental setup, PSTVd also caused more severe symptoms and accumulated to high levels (Fig. 1A to 1D) when infected tomato plants were grown at a higher temperature.

**Figure 1.**
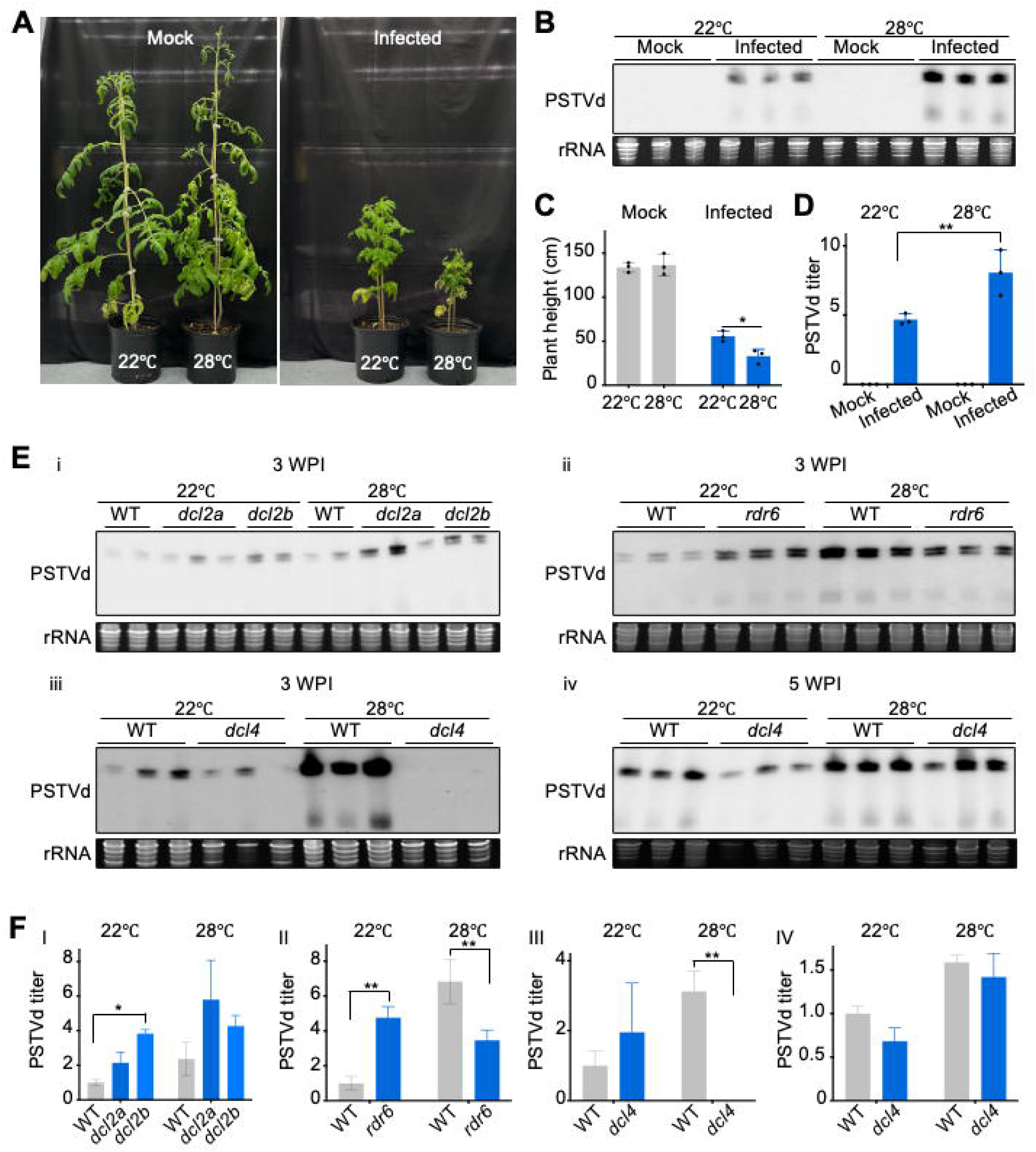
Temperature modulates PSTVd infectivity and the antiviroidal activity of RNA silencing components. (**A**) PSTVd-infected plants had severer stunting symptoms at 28 °C. (**B**) PSTVd titer was significantly higher when plants were grown at 28 °C. rRNAs serve as the loading control. Normalized data were subject to t-test using Prism. (**C**) PSTVd titers in monogenic mutant tomato plants (*dcl2a*, *dcl2b*, *dcl4*, and *rdr6*) grown in 22 °C or 28 °C. Ethidium bromide staining of rRNAs served as loading control. (**D**) Quantified data were analyzed using t-test by Prism. Only statistically different pairs are marked. *, <0.05; **, <0.01.

We first analyzed whether host RNA silencing components play a role in modulating PSTVd titers at different temperatures. We obtained established tomato monogenic mutant lines and performed infection analyses. In *dcl2b* loss-of-function tomato plants (*16, 34*), PSTVd titer was significantly higher than in the control plants when growing at 22°C but the differences were less pronounced when plants were grown at 28°C (Fig. 1Ei and 1F). In the established tomato *rdr6* mutant (*35, 36*), PSTVd titer was significantly higher than in the control wild-type tomato when grown at 22°C but was significantly lower than that in the control plants when grown at 28°C (Fig. 1Eii and 1F). Interestingly, in the *dcl4* plants (*35, 36*), PSTVd had a significantly delayed infection at 28°C but not at 22°C (Fig. 1Eiii, Eiv and 1F). Overall, the antiviroidal activity of SlDCL2 genes was impaired at 28°C, but RDR6 and DCL4 appeared to become pro-viroid at 28°C. Therefore, analyzing the PSTVd infectivity differences at 22°C and 28°C can provide new insights into host defense mechanism against viroids, an unusual group of pathogens that are composed of circular noncoding RNAs.

### Elevated temperature promotes a more uniform RNA conformation in favor of infection

Given that temperature affects the thermodynamic folding of RNAs (*37*) and that RNA structural motifs are critical for viroid infection (*38*), we aimed to analyze the PSTVd RNA structurome in tomato and determine whether RNA folding also contributes to the infectivity. High-quality nanopore-based *in vivo* RNA structure profiling data (smStructure-seq) were obtained for analyzing PSTVd conformations in planta (Fig. S1). This new technology offers single-molecule resolution for understanding the folding of RNA populations in cells. Global comparison of dimethyl sulfate (DMS) reactivity revealed a clear temperature-dependent structural shift. The distribution of DMS reactivities and the overall DMS reactivities showed a significant increase at 22°C compared to 28°C (Fig. 2A), indicating that the PSTVd RNAs adopt more single-stranded and flexible conformations at 22°C. Shannon entropy, which reflects the structural heterogeneity of RNA, had a higher value at 22°C, indicating increased conformational flexibility and the presence of multiple alternative structures (Fig. 2B). The Gini index, which measures the overall unevenness of DMS reactivity distribution along the RNA and is widely used as an indicator of RNA structural organization, was significantly elevated at 28°C (Fig. 2B), reflecting an overall uneven and highly structured RNA conformation.

**Figure 2.**
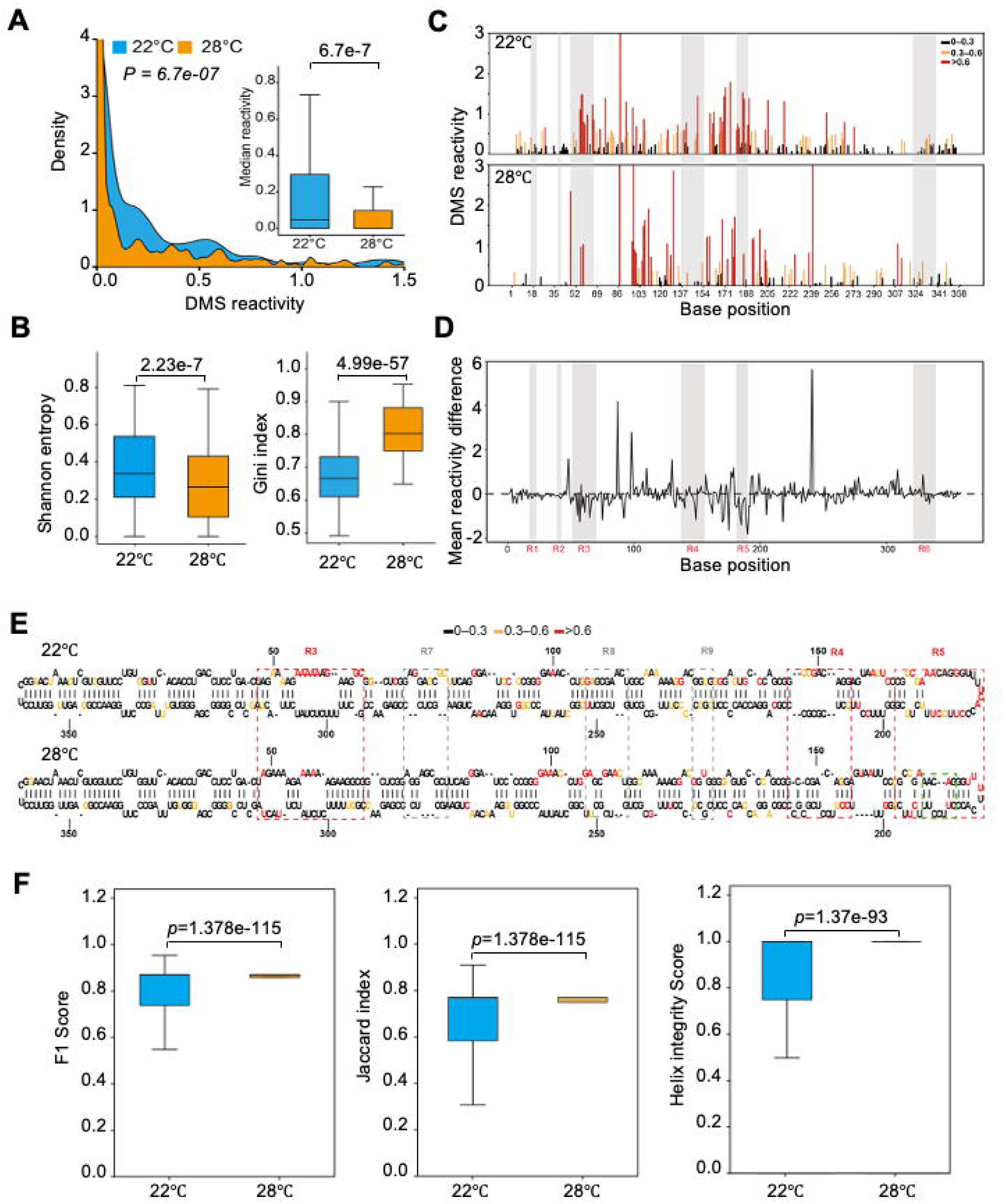
Temperature-dependent folding of PSTVd RNA revealed by DMS reactivity profiling. (**A**) Distribution of DMS reactivities at 22°C (blue) and 28°C (orange). Statistical significance was assessed using the Wilcoxon rank-sum test (P = 6.78 × 10⁻□). Inset boxplot displays median reactivity differences between the two conditions (Mann–Whitney U test). (**B**) Distribution of median Shannon entropies (left) and Gini indexes (right) under 22°C and 28°C conditions. In all box plots, boxes represent the interquartile range (25th–75th percentiles), and the central line indicates the median. Whiskers extend to 1.5 × IQR. Outliers (defined as values beyond 1.5 × IQR) are not displayed. Statistical significance was assessed using a two-sided Mann–Whitney U test. (**C**) Per-nucleotide DMS reactivity profiles of the PSTVd under 22°C and 28°C conditions. Reactivity values are color-coded (black: 0–0.3; orange: 0.3–0.6; red: >0.6). Shaded regions indicate regions with significantly different reactivity identified by DiffScan analysis. (**D**) Mean DMS reactivity differences (28°C vs 22°C) across the PSTVd genome are shown along nucleotide positions. Positive values indicate increased reactivity (reduced structural constraint) at 28°C, whereas negative values indicate higher reactivity at 22°C. Shaded regions represent segments with statistically significant differences identified by DiffScan analysis. Six distinct differential regions (R1–R6) are highlighted. (**E**) Secondary structure models of PSTVd at 22°C (top) and 28°C (bottom), with DMS reactivity overlaid. Regions R3–R5 (dashed red boxes) exhibit clear structural rearrangements between temperatures. Dashed gray boxes indicate additional regions showing apparent structural changes, although they were not identified as statistically significant by DiffScan analysis. C-loop in the R5 region (28°C) is highlighted by a green box. (**F**) Distribution of structural similarity scores at the single-molecule level under 28°C and 22°C. Boxplots show F1 score, Jaccard index, and helix integrity score, respectively. Center lines indicate medians, boxes represent the interquartile range (IQR), and whiskers extend to 1.5 × IQR. Statistical significance between conditions was assessed using the Mann–Whitney U test, and exact P values are indicated above each comparison.

Per-nucleotide DMS reactivity profiles further revealed substantial heterogeneity along the PSTVd genome under both temperature conditions. At 22°C, elevated reactivity signals were observed at multiple positions, indicating the presence of locally flexible or single-stranded regions (Fig. 2C). By contrast, the reactivity pattern at 28°C showed a redistribution of highly reactive sites, with some positions displaying increased reactivity while others exhibited reduced signals (Fig. 2C), indicating the structural differences between 22°C and 28°C conditions, consistent with the global analyses of Shannon entropy and Gini index. Using DiffScan (*39*) to detect local differences in reactivity, we identified six differential regions (R1–R6) between the two conditions (Fig. 2D), indicating localized temperature-sensitive structural changes. Furthermore, DMS reactivity–constrained RNA fold analysis also revealed multiple regions with apparent structural differences between 22°C and 28°C, including R3, R4, and R5 that overlapped with the DiffScan-identified regions (Fig. 2E). These observations supported the robustness of these temperature-responsive structural changes. Using three distinct statistical analyses (Fig. 2F), we found that the R5 region showed a wide range of alternative conformations at 22°C, while its conformation was uniform at 28°C. The dominant conformation at 28°C possesses the essential C-loop, which is required for nearly all nuclear-replicating viroids(*12*). The C-loop is involved in binding to VirP1 and utilizing IMPA4-based machinery for the nuclear entry, and even multiple isosteric point-mutations within the C-loop impair VirP1 binding and reduce nuclear import efficiency (Fig. S2), indicating the importance of the C-loop for viroid infectivity (*12*). Therefore, the poor replication of PSTVd at lower temperatures may be associated with the presence of only a small fraction of PSTVd RNAs that form the C-loop structure.

### NPR1 exerts anti-PSTVd activity at cool ambient temperature

A previous study showed that citrus exocortis viroid (CEVd), another nuclear-replicating viroid, establishes systemic infection earlier and elicits stronger symptoms in *NahG*-overexpression (OE) tomato plants, in which salicylic acid is degraded by the *NahG* gene (*40*). Interestingly, salicylic acid (SA) signaling and the SA receptor NPR1 both function in a temperature-dependent manner (*41*). By using an established tomato *NPR1*-OE line (*42*), we found that the PSTVd titer was significantly lower in the *NPR1*-OE tomato at 22°C but not at 28°C (Fig. 3A), indicating that NPR1 also participates in repressing PSTVd titer in tomato, and the efficacy is reduced at higher temperature. By using virus-induced gene silencing, we found that *NPR1* silencing led to higher PSTVd accumulation in tomato plants (Fig. 3B). Therefore, these data demonstrated that NPR1 possesses an antiviroidal activity, but the activity is insufficient at high ambient temperature.

**Figure 3.**
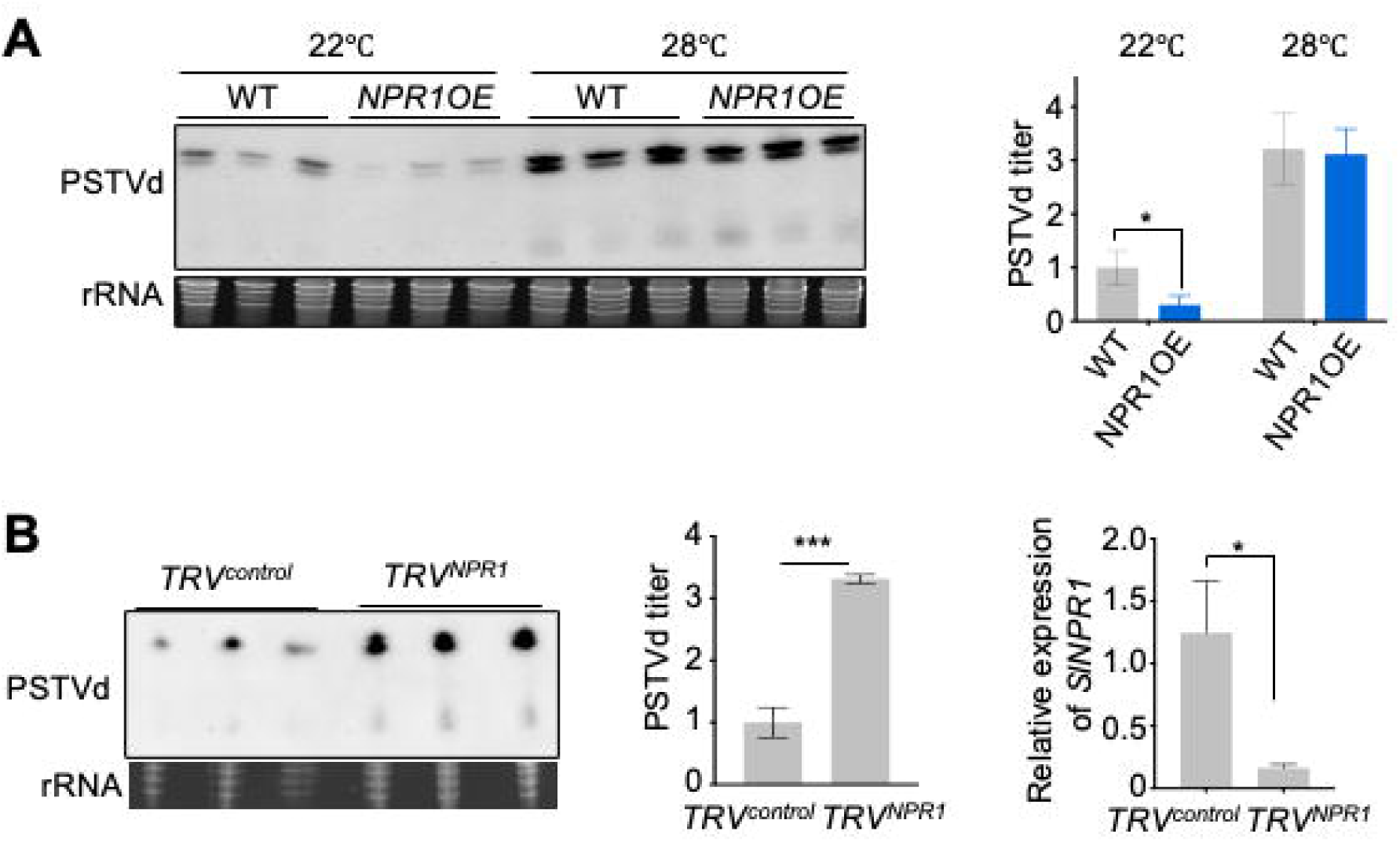
NPR1 identified as a new resistance gene against PSTVd in tomato. (**A**) RNA gel blots showed PSTVd titers in NPR1-overexpression (OE) tomato. Ethidium bromide staining of rRNAs served as loading control. (**B**) Tobacco rattle virus (TRV)-based gene silencing of *NPR1* led to increased PSTVd titer at 22°C. RT-qPCR showed the silencing effect of *NPR1*. Quantified data were subject to t-test. Only statistically different pairs are marked. *, <0.05; ***, <0.001.

Since the nuclear accumulation of NPR1 is often triggered by SA, we asked whether PSTVd infection induces the level of SA. Using HPLC, we measured SA content in PSTVd-infected tomato plants. The data showed that PSTVd infection significantly induced the accumulation of free SA and total SA (Fig. 4A). The level of salicylate glucoside (SAG), a major storage and inactive form of SA, also showed the trend of increase (Fig. 4A). The data demonstrate the activation of SA synthesis triggered by PSTVd infection.

**Figure 4.**
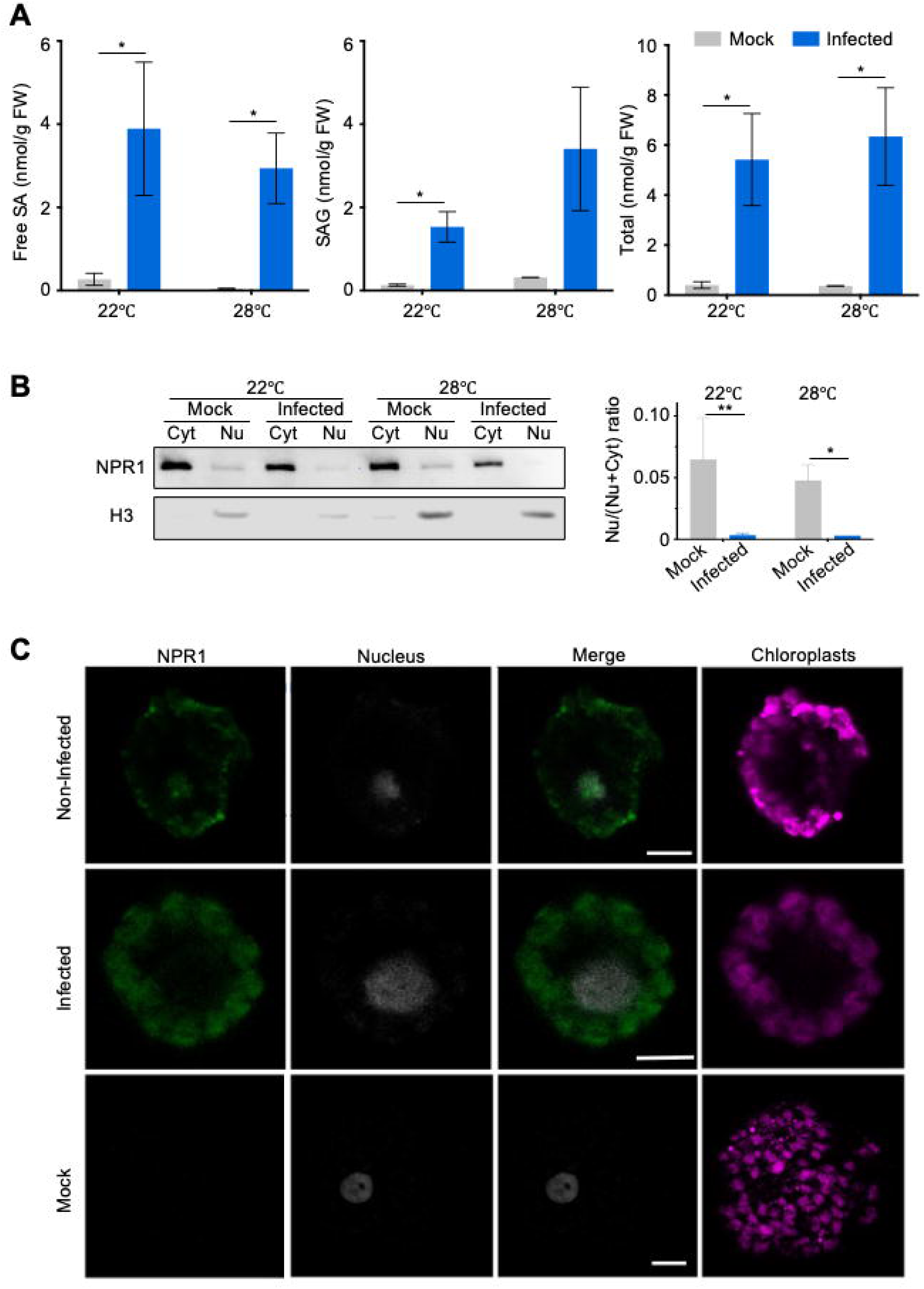
NPR1 failed to accumulate in PSTVd-infected nuclei despite the induction of salicylic acid. (**A**) PSTVd infection triggered the induction of active form salicylic acid (SA) was well as the storage form (SAG), as illustrated by HPLC analysis. Infection was confirmed by RNA gel blots (Fig. S3) (**B**) immunoblots and (**C**) confocal microscope observations showed that nuclear accumulation of NPR1-GFP was impaired in PSTVd-infected protoplasts. Nu, nuclear fraction. Cyt, cytoplastic fraction. H3, Histone H3. GAPDH served as loading control. N and P represent mock and infected plants, correspondingly. DAPI staining marks the nucleus. Scale bar, 5μm. Quantified data were subject to t-test. Only statistically different pairs are marked. *, <0.05; **, <0.01.

### NPR1 nuclear accumulation was impaired in PSTVd-infected plants at higher temperatures

To understand why NPR1 exhibited weaker antiviroidal activity at 28°C despite increased SA levels, we analyzed the subcellular distribution of NPR1 in mock and PSTVd-infected *N. benthamiana* protoplasts. Using immunoblots to quantitatively analyze the NPR1 accumulation in the cytoplasmic and nuclear fractions, we found that the nuclear accumulation of NPR1 was significantly impaired in the infected protoplasts, particularly at 28°C (Fig. 4B). Confocal microscopy analysis also confirmed the observation (Fig. 4C).

To understand why NPR1 failed to accumulate in nuclei at 28°C despite the PSTVd-induced increase in SA levels, we performed non-targeted proteomics analyses to investigate the underlying mechanism. We found a significant reduction of Nup88 and IMPA3, which are the key factors in regulating NPR1 nuclear accumulation (*43, 44*). In fact, there was a systematic reduction of multiple nucleoporins (GP210, Nup85, Nup88, Nup96, Nup133, Nup155, Nup205) and nuclear transport receptors (IMPA1, IMPA2, IMPA3, IMPA9, IMPB1-1, IMPB1-2, IMPB3, SAD2) (Fig. 5A to 5D)(Dataset 1 and 2). Testing of selected Nup and IMPA proteins using transient expression in mock or PSTVd-infected *N. benthamiana* also confirmed the observation by proteomics (Fig. 5E).

**Figure 5.**
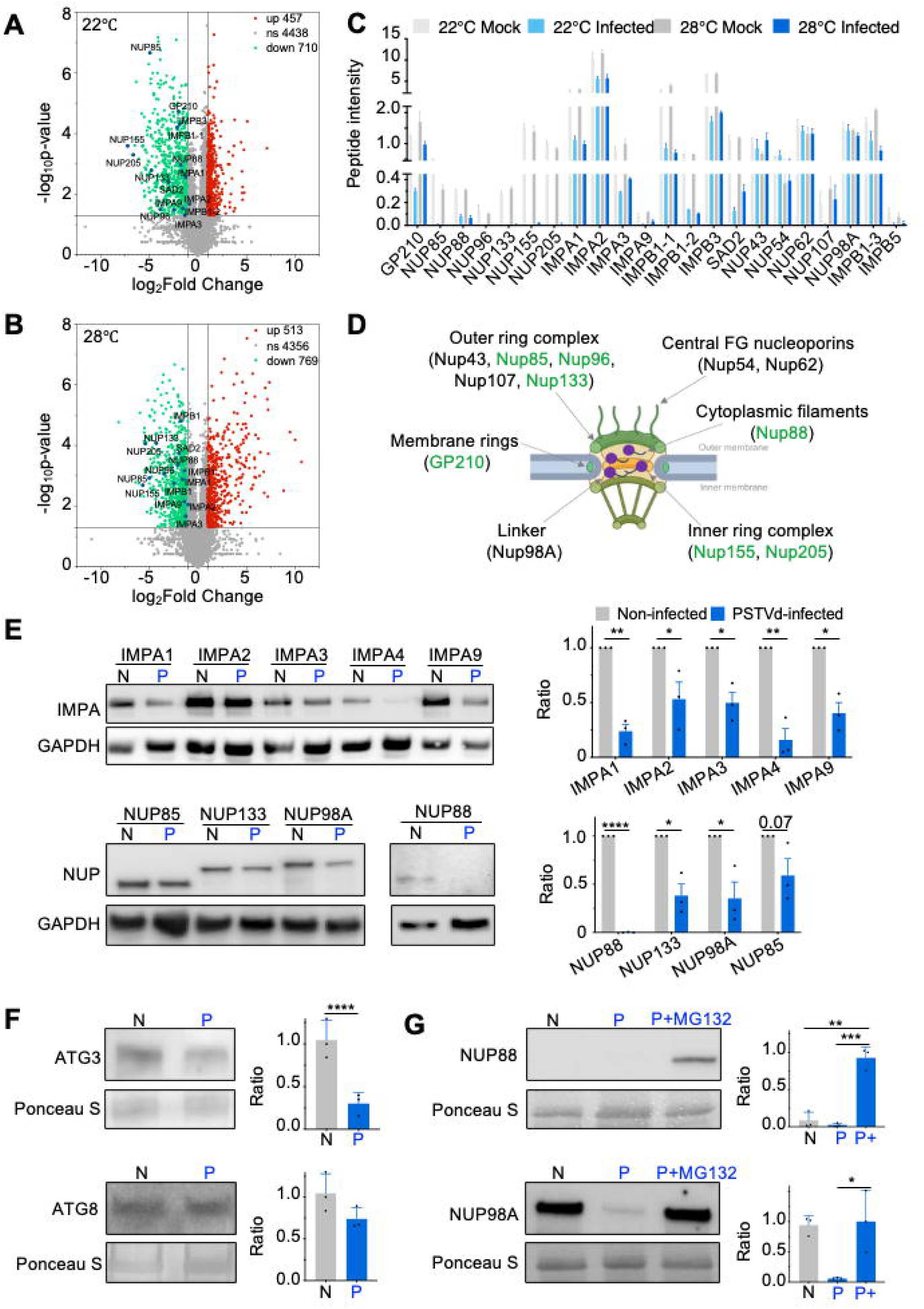
Repressed accumulation of nucleoporins and importins. (**A**) and (**B**) Volcano plots and (**C**) column graph showed the protein abundance of nucleoporins and importins detected by nontargeted label-free proteomics from mock and PSTVd-infected tomato leaves growing at 22°C and 28°C. Infection was confirmed by RNA gel blots (Fig. S4). Notably, these genes’ mRNA levels showed a distinct pattern (Dataset S2). (**D**) A nuclear pore model illustrates the position of nucleoporins. Reduced protein factors are highlighted in green. Most of reduced nucleoporins mainly reside in the nuclear pore complex (NPC) core scaffold (inner, outer, and membrane) and in the cytoplasmic filaments, implying a potential reorganization of the NPC. Certain nucleoporins were not affected, implying that the repression is selective. (**E**) Nucleoporins and importins accumulated to lower levels in PSTVd-infected *N. benthamiana* in agroinfiltration assay, as compared with expression in mock plants. GAPDH served as internal control. (**F**) Immunoblotting showed that autophagy marker, ATG3, was reduced in infected plants. ATG8 was not affected. This observation suggests that autophagy pathway is not involved in this repressed accumulation of nucleoporins and importins. (**G**) MG132 treatment restored the accumulation of Nup88 and Nup98A in transient expression assay, indicating that 26S proteasome activity is induced upon PSTVd infection and leads to the repression of Nup proteins. Ponceau S staining of rubisco served as loading control. Quantified data were subject to t-test. Only statistically different pairs are marked. *, <0.05; **, <0.01; ***, <0.001; ****, <0.0001.

Since the transcripts level of most nucleoporins and nuclear transport receptors remained unchanged (Dataset 2), the reduced accumulation of these proteins is likely attributed to post-translational regulation. Nucleoporins may be degraded through the autophagy pathway (*45, 46*) or the 26S proteasome pathway (*47*). We first tested whether the autophagy pathway was activated upon PSTVd infection by detecting the ATG8, which is involved in recruiting cargo into the autophagosomes (*48*), and ATG3, whose activity promotes the ATG8 autophagosome elongation(*49*). Interestingly, endogenous ATG3 was significantly reduced in the PSTVd-infected plants (Fig. 5F), suggesting the repression of autophagy activity based on the current model (*50, 51*). On the other hand, we tested the effect of MG132 on nucleoporin accumulation. The protein levels of Nup88 and Nup98a were significantly induced after MG132 treatment in the infected leaves (Fig. 5G), supporting that the reduced accumulation of nucleoporins is likely caused by induced 26S proteasome activity.

Given the large-scale repression of nucleoporins and nuclear import adapters, we reasoned that additional SA level would not improve NPR1 nuclear accumulation. In support of this assumption, we found that NPR1 nuclear accumulation was not improved in PSTVd-infected cells after the supplementation of acibenzolar-S-methyl (abbreviated as ASM or BTH, a salicylic acid analog), which increased NPR1 nuclear accumulation in noninfected cells (Fig. 6A). On the other hand, we assume that blocking the 26S proteasome activity may restore the level of nucleoporins and nuclear import adapters, which may improve NPR1 nuclear accumulation. As shown in Fig. 6B, NPR1 could accumulate in the PSTVd-infected nuclei after a 4 hr incubation with MG132, supporting the importance of the nucleoporins and nuclear import adapters in NPR1 nuclear accumulation.

**Figure 6.**
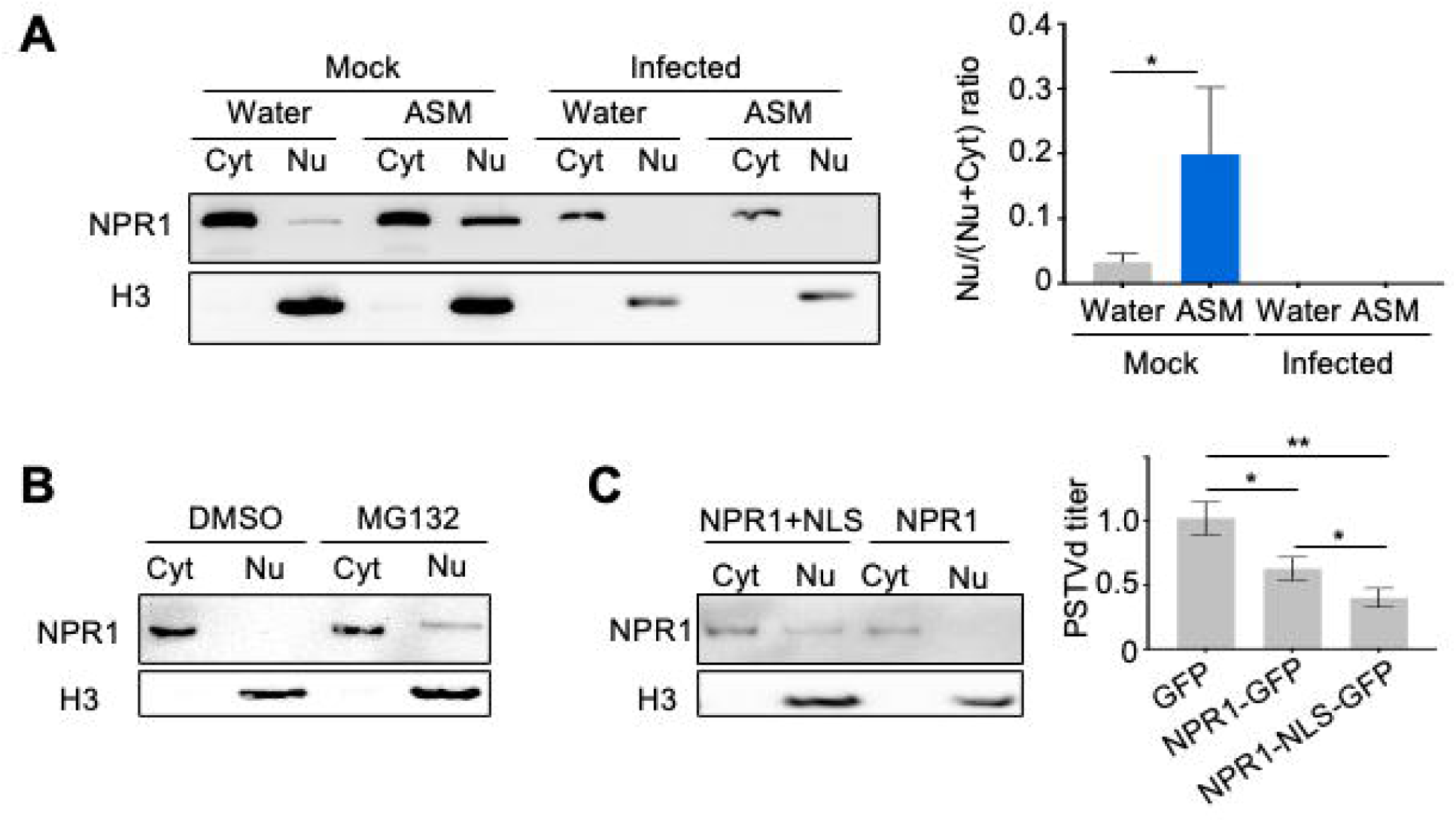
NPR1-based antiviroidal activity is impaired due to repressed accumulation of nucleoporins and importins. (**A**) Increasing SA content induced NPR1 nuclear accumulation in mock but not PSTVd-infected protoplasts. Immunoblots demonstrated the accumulation levels of NPR1. Nu, nuclear fraction. Cyt, cytoplastic fraction. H3, Histone H3 (nuclear marker). (**B**) MG132, which could restore the accumulation of nucleoporins, increases NPR1 nuclear accumulation. (**C**) Fusing a Simian virus 40 large T-antigen nuclear localization signal (NLS) to NPR1 to increase its nuclear localization, which led to reduced PSTVd titers. RT-qPCR was used to quantify PSTVd titers, using 18S rRNA for normalization. Quantified data were subject to t-test. Only statistically different pairs are marked. *, <0.05; **, <0.01.

To directly link the antiviroidal activity of NPR1 and its nuclear accumulation, we introduced a well-established simian virus 40 (SV40) nuclear localization signal (NLS) between NPR1 and C-terminally-fused GFP as previously described (Fig. S5)(*52*). Not surprisingly, SV40 NLS only partially increased NPR1 nuclear accumulation (Fig. 6C), reflecting the repression of nucleoporins and nuclear transport adapters. Since NPR1 retains antiviroidal activity at 22°C but not 28°C, we transformed PSTVd-infected *N. benthamiana* protoplasts and incubated at 22°C. RT-qPCR analysis showed that PSTVd titer was further reduced when NPR1 accumulated more in the nucleus with the help of SV40 NLS, supporting that NPR1 antiviroidal activity relies on its nuclear accumulation.

### PSTVd expands its host range to *Arabidopsis* at high ambient temperatures

*A. thaliana* is a known nonhost of viroids. Despite that PSTVd can replicate in *Arabidopsis* protoplasts, our previous trial of infecting *A. thaliana* grown at 22°C was not successful (*53*). Given that PSTVd has a significantly higher infectivity in tomato at 28°C, we tried to infect *A. thaliana* plants growing at 28°C. Interestingly, PSTVd progeny was detected in the RNA samples pooled from the wild-type (WT) and *npr1-3* plants (Fig. 7A). However, no noticeable symptoms were observed. In the WT plants, only very few stochastic mutations occurred in addition to the majority sequence of inoculum (Fig. 7B). By contrast, the progeny sequences in the *npr1* plants are uniform as Int^G201U^, which has a G at the 201 position as compared with the standard lab strain Int (Fig. 7B and 7C). This correlated with a higher PSTVd titer in the *npr1-3* plants. We named this new Int^G201U^ strain, ‘Biao Ding (BD) strain’, in honor of this researcher’s contributions to the viroid research. The BD strain was infectious and genetically stable in *A. thaliana* (Fig. 7D), *N. benthamiana* and tomato.

**Figure 7.**
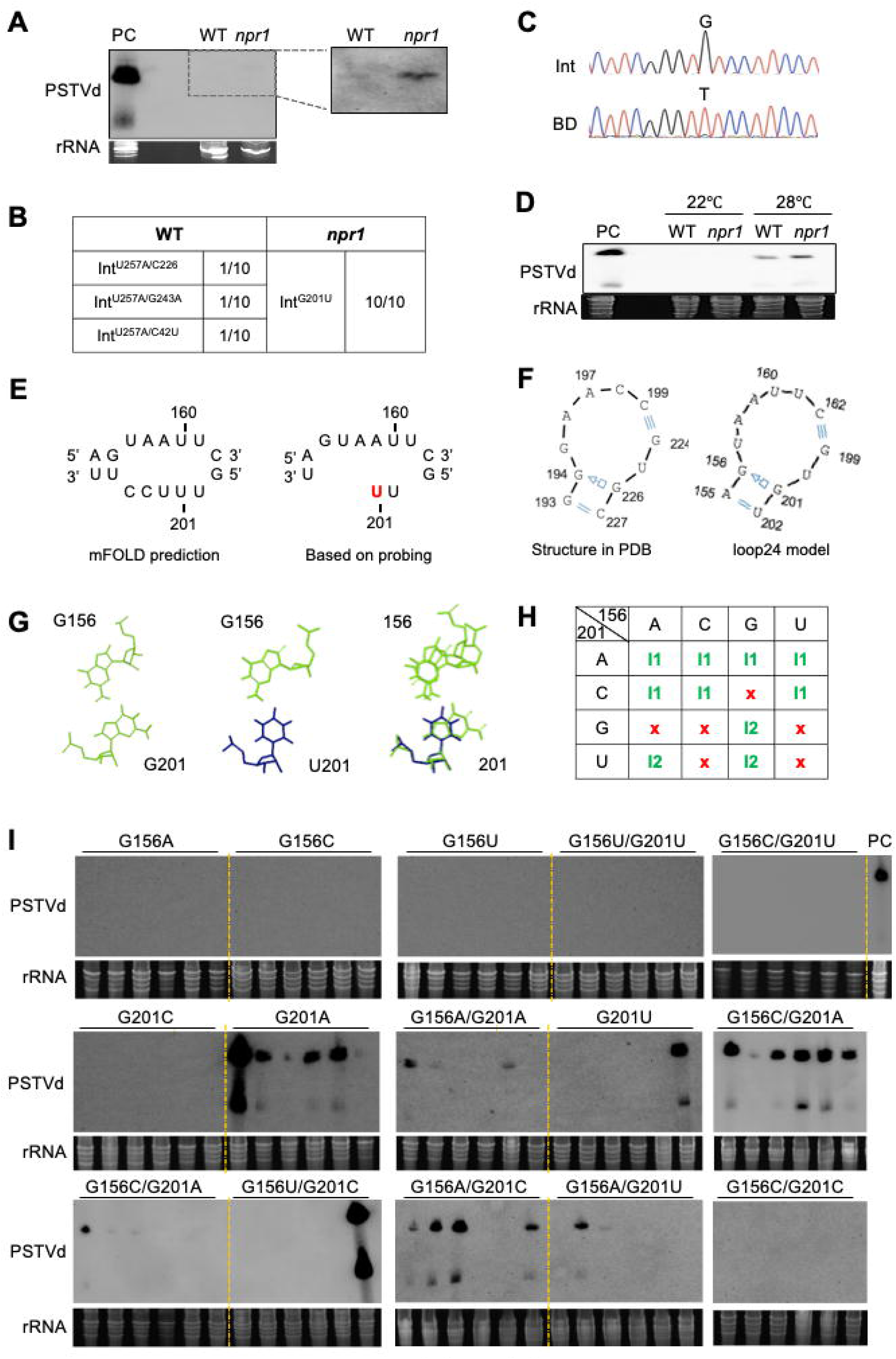
**PSTVd-infection in Arabidopsis led to the emergence of the G201U mutation**. (**A-C**) PSTVd could systemically infect *Arabidopsis* wildtype (WT) and the *npr1-3* mutant, shown after over-exposure of RNA gel blots. The higher PSTVd titer in the *npr1-3* mutant led to the emergence of Int^G201U^. (**D**) Inoculation using Int^G201U^ led to *Arabidopsis* infection. In WT, the successful rate was 2 out of 8. In the *npr1-3* mutant, the successful rate was 4 out of 8. (**E**) Two versions of loop 24 secondary structures based on SHAPE (left) and multiple probing (right). (**F**) Structural modeling of the loop24. Blue triangles depict the sugar edge. Blue boxes depict the Hoogsteen edge. SRP, signal recognition particle. (**G**) Structural views of G-G (left) and G-U (middle) trans-Hoogsteen-Sugar pairings, as well as the overlay (right), were shown. (**H**) The table summarized RNA base catalog data regarding the isosteric groups (I1 and I2). I1 and I2 only differ slightly in C1’-C1’ distances. The red crosses indicate that the base pairings cannot form between the two nucleotides. (**I**) Non-isosteric mutants (labeled in purple color) failed to infect tomato plants, while isosteric variants (labeled in green color) retain the infectivity. Ethidium bromide staining of rRNAs served as loading control. PC, positive control.

The nucleotide at the 201 position resides in the PSTVd loop 24. Although it is often described as the “UAAUU…UUUCC” motif (Fig. 7E), a JAR3D search (*54*) using this sequence as a query against experimentally validated RNA structures in the Protein Data Bank (PDB) did not identify any high-confidence models. Notably, multiple chemical probing analyses (*55*) (RNase S1 digestion, RNase T1 digestion, RNase A digestion, Dimethyl Sulfate probing, and partial cleavage by heavy metals like Pb^2+^ and Tb^3+^) (Fig. S6) suggested that this local structure could fold as “GUAAUU…UG” (Fig. 7E). Using this revised structure as a query identified only one high-confidence RNA structural model. This model is based on a motif in the Signal Recognition Particle (SRP) RNA from *Sulfolobus solfataricus* (PDB id: 3KTW) (Fig. 7F). This motif in SRP RNA is not directly involved in protein binding but resides between the binding sites of two proteins, SRP19 and SRP54 (*56*). Based on this model, only the G156 uses its sugar edge to pair with the Hoogsteen edge of G201 to form a trans-Sugar-Hoogsteen pairing, and other nucleotides in the loop are not involved in pairing (Fig. 7F).

Based on the listing in RNA Basepair Catalog (*57*), G201 and U201 are isosteric (Fig. 7G), which means that they may be functionally interchangeable by sharing similar C1’-C1’ distance, the same edge-edge interactions and the same orientation of glycosidic bonds (*58*). We then tested all 10 isosteric variants (including WT as a control and G201U)( Fig. 7H) and all six non-isosteric variants to genetically test this model via analyzing their infectivity. As expected, none of the non-isosteric variants was infectious, whereas nearly all isosteric variants remained infectious in tomato except Int^G156C/G201C^ (Fig. 7I). The exception of Int^G156C/G201C^ might be due to other selection pressure, and similar exceptions have been reported previously (*59, 60*). Overall, the functional mutagenesis validated that either G (in the Int strain) or U (in the BD strain) at the 201 position uses the Hoogsteen edge to pair with the sugar edge of G at the 156 position.

### Salicylic acid analog can block PSTVd infection

Although NPR1 failed to accumulate in the nuclei of PSTVd infected cells even after exogenous application of ASM, ASM treatment has been reported to delay CEVd systemic spreading in tomato (*40*). This discrepancy implies that excessive SA may function independently of NPR1 to inhibit the infection of nuclear-replicating viroids, which may be exploited for chemical control of viroid infection. To explore this possibility, we first compared the efficacy of the reported viroid inhibitors, echinocystic acid (EA)(*61*) and ASM(*40*). We found a striking effect of ASM in completely blocking PSTVd infection in ∼75% plants, a significant improvement over mock-treated plants (100% infected) or plants sprayed with EA (100% infected)(Fig. 8A). It is noteworthy that it was achieved by spraying ASM solution daily to PSTVd-inoculated tomato seedlings in the first week, which increased the spraying times as compared with the previous report (*40*).

**Figure 8.**
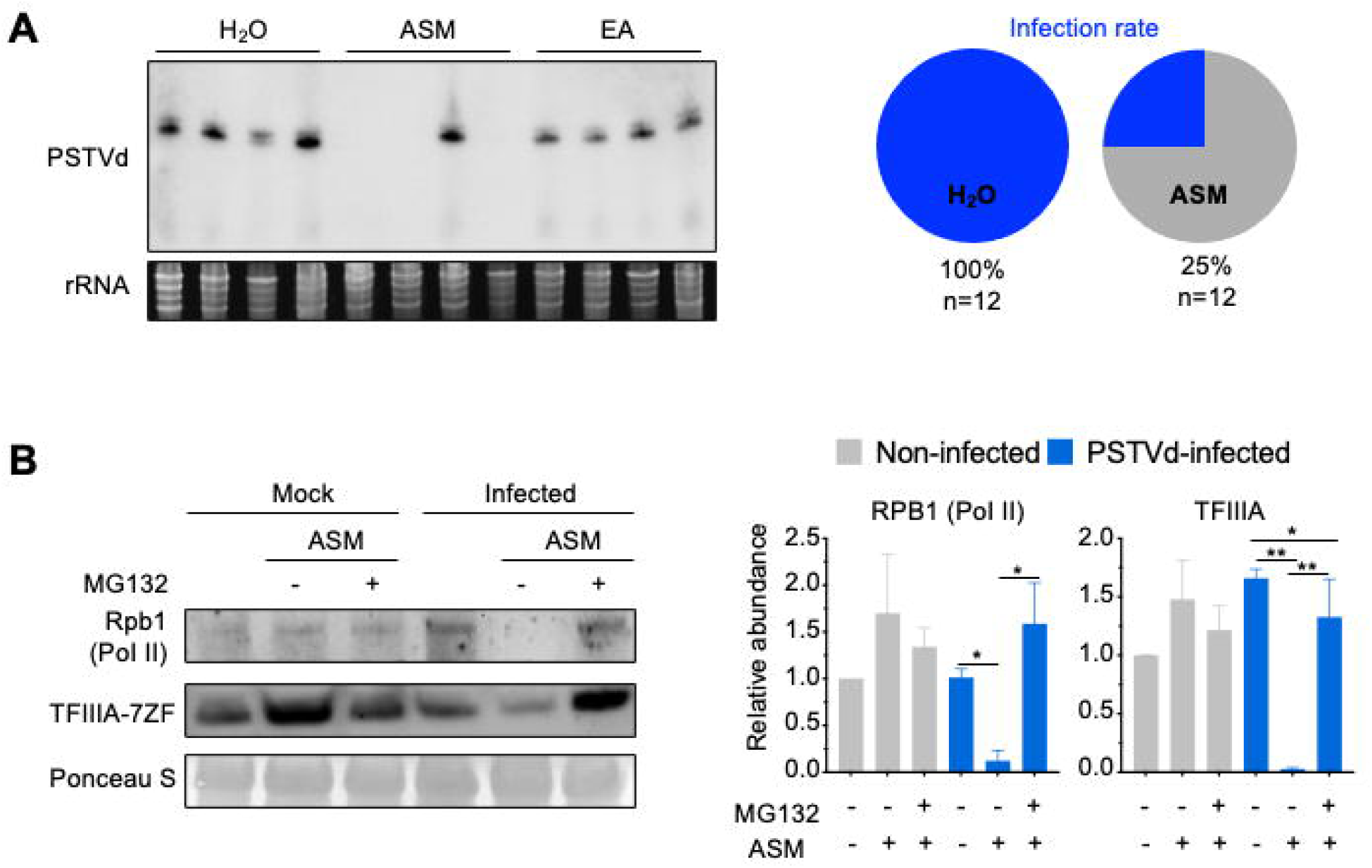
Chemical inhibition of PSTVd infection. (**A**) Spraying acibenzolar-S-methyl (ASM) blocked PSTVd infection in 75% of inoculated plants. Ethidium bromide staining of rRNAs served as loading control. EA, echinocystic acid. (**B**) Spraying ASM reduced Rpb1 and TFIIIA-7ZF only in PSTVd-infected plants but not in mock plants. MG132 rescued protein level. Ponceau S staining of rubisco served as loading control. Quantified data were analyzed using t-test. Only statistically different pairs are marked. **, <0.01; ****, <0.0001.

As showed in Fig. 6A, this supplement of ASM cannot restore NPR1 nuclear accumulation. To understand a possible mechanism underlying this inhibition, we tested the endogenous expression levels of key host factors, Pol II and TFIIIA-7ZF. Pol II. In particular, the largest subunit of Pol II (Rpb1) is known to be degraded through 26S proteasome activity under stress conditions (*62, 63*). Therefore, we included a 26S proteasome inhibitor, MG132, in the treatment. Both Rpb1 and TFIIIA-7ZF were significantly reduced in PSTVd-infected samples after the ASM spray (Fig. 8B). MG132 treatment could prevent the reduction, indicating that 26S proteasome activity was involved in the degradation of both proteins during PSTVd infection.

## Discussion

Elevated temperature clearly enhanced PSTVd infectivity in tomato plants (Fig. 1). This effect is partly attributed to the reduced defense activity of DCL2b at 28°C. Some RNA silencing components, such as RDR6 and DCL4, also contributed to the phenomenon through acting like pro-PSTVd factors at 28°C (Fig. 1E and 1F). The data demonstrated that RNA silencing does not confer overall antiviroidal activity at 28°C. Interestingly, *in vivo* smStructure-Seq analysis demonstrated that PSTVd adopted a more uniform structure *in vivo* in favoring infection at 28°C, which contrasts with the presence of multiple alternative folding conformations at 22°C. In the relatively uniform conformation at 28°C, the C-loop critical for PSTVd nuclear entry (*12*) was apparent and dominant, which explains the higher replication efficiency of PSTVd under this condition (Fig. S2). Notably, this is the first *in vivo* RNA structurome analyses of a plant pathogenic RNA in host plants. Compared with previous analyses that only aggregate probing data to deduce one overall structure, smStructure-Seq and ensemble analyses outlined the individual structures in the population, thereby helping understand the structural folding dynamics of viroid RNA genomes with more details.

In addition, we identified NPR1 as a novel antiviroidal factor, whose activity is also impaired by elevated temperature (Fig. 3A). In PSTVd-infected cells at 28°C, NPR1 did not accumulate in the nucleus (Fig. 4B and 4C), despite the SA level was induced by PSTVd (Fig. 4A). This correlated with massive reductions in nucleoporins and nuclear transport receptors, including Nup88 and IMPA3 that have been reported to regulate NPR1 nuclear accumulation (*43, 44*). The repressed accumulation of Nup and IMPA proteins is likely attributed to induced 26S proteasome activity (Fig. 5G). In line with the observation, activation of SA signaling by exogenously applied ASM did not force NPR1 to accumulate in the nucleus in infected cells (Fig. 6A), supporting that the nuclear transport adapters and/or the nuclear pore complex are compromised therein. By contrast, treating cells with MG132, which is thought to restore the shuttling machinery and the nuclear pore complex by blocking proteasome activity, could restore NPR1’s nuclear accumulation in PSTVd-infected nuclei (Fig. 6B). Together, these findings expand the defense role of NPR1 to a new class of pathogens, which are merely pieces of circular noncoding RNAs that do not encode any proteins. Importantly, the data clearly establish that plant immune factors can pose negative effects on viroid infection. Our finding also illustrates a new mechanism that impairs the defense activity of NPR1 through repressing the large portion of nucleoporins and importin factors. Although many eukaryotic pathogens tend to manipulate the nuclear pore complexes and the nuclear transport receptors to facilitate their infection(*64, 65*), such a wide range of reduction in the nuclear transport receptors and nucleoporins was not previously documented.

Based on the knowledge of NPR1’s antiviroidal function and increased PSTVd infectivity at elevated temperature, we attempted to infect *A. thaliana* with PSTVd at 28°C, using both WT and *npr1-3* mutant (Fig. 7A). PSTVd could only infect *A. thaliana* at 28°C but not 22°C, and accumulated to a much higher titer in the *npr1-3* mutant. This high level of PSTVd titer allowed the selection of the optimal variant Int^G201U^. Through motif modeling and functional mutagenesis analysis, we demonstrated that G201 and U201 are isosteric, retaining the same edge-edge interaction pattern, similar C1’-C1’ distances, and the same orientation of glycosidic bonds. Despite being an isosteric mutation, U201 appears to be strongly selected in both dahlia and *A. thaliana*, whereas G201 is dominant in tomato and potato (*66–68*). Dahlia is an emerging non-solanaceous host of PSTVd, and all reported PSTVd strains in dahlia have U at the 201 position (Table S1). Infected dahlia plants in the field could also serve as reservoirs for transmission to susceptible plants (i.e., tomato) in the surrounding areas (*29*), posing a significant threat to crop production and safety. Nevertheless, strengthening the SA-based immunity may still help combat the infection of nuclear-replicating viroids in the future.

Notably, exogenously supplying ASM did not allow NPR1 to accumulate in the infected nuclei (Fig. 6A) but still blocked PSTVd infection in most plants (Fig. 8A). This repression activity is likely through degrading the host factors necessary for viroid replication. Although TFIIIA-7ZF and Pol II appear to exhibit similar expression levels between the mock and infected plants, spraying ASM broke the balance and led to the degradation of both proteins (Fig. 8B). Intriguingly, spraying ASM onto mock plants did not lead to Pol II degradation. This may be attributed to the Pol II remodeling for PSTVd RNA-templated transcription. During replication, the 12-subunit Pol II is reorganized to a 7-subunit complex (containing Rpb1,2,3,8,10,11,12)(*15, 16*). This remodeling, particularly the lack of Rpb4,6,7, may be recognized as an assembly deficiency, leading to Rpb1 degradation (*69*). Overall, this funding confirms that chemical control of PSTVd infection is possible.

Based on the data, we propose a model that temperature modulates PSTVd infectivity through promoting the uniform folding of PSTVd genome RNA and repressing the antiviroidal activity of RNA silencing and NPR1 (Fig. 9). This increased infectivity at high temperature permits PSTVd to jump to new hosts, which is a possible scenario during the prolonged high temperatures in the real world with ongoing global climate changes. The message highlights a pressing high risk of outbreaks of new viroid diseases. Our data also suggest that engineering thermo-tolerable RNA silencing and NPR1 activities may be the direction for molecular breeding in combating future PSTVd outbreaks. SA-based chemical treatment can also be exploited to block infection in crops.

**Figure 9.**
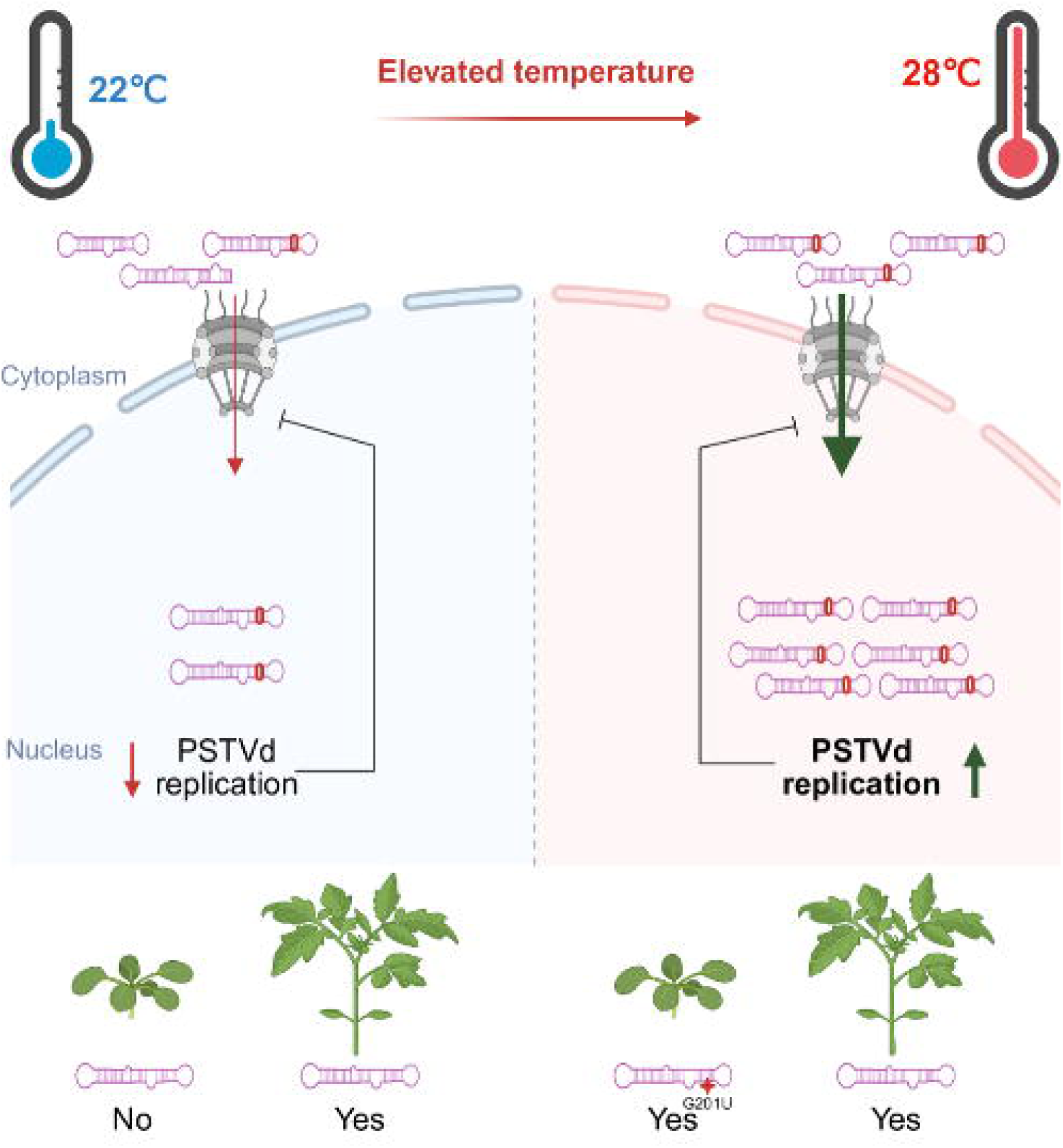
The model of distinct climate conditions modulating PSTVd infectivity and host range. The nuclear entry signal, C-loop, is depicted by bold red circles in RNA molecules.

## Materials and Methods

### Plant material and growth condition

Tomato (*Solanum lycopersicum*) and *N. benthamiana* were germinated in the dark, transferred into soil, and grown under a 14 h light/10 h dark photoperiod at 22°C or 28°C. The *A. thaliana* plants were germinated on ½ MS plate and transferred to ½ vermiculite and ½ pot soil with a 14 h light/10 h dark photoperiod at 22°C. After viroid inoculation, *A. thaliana* plants were kept at 22°C or 28°C as described in the results. The previously described T-DNA plasmid pBI1.4T-AtNPR1 (*70*) was used to transform the tomato cultivar ‘Money Maker’ following an *Agrobacterium tumefaciens*-mediated genetic transformation protocol (*42*). The tomato genetic transformation experiment was conducted by the University of Nebraska-Lincoln Plant Transformation Facility. Various tomato monogenic mutant lines were obtained from Tomato Genetics Resource Center at University of California, Davis. Tomato *dcl2a* and *dcl2b* loss-of-function mutants generated via Crispr-technology was gift from Dr. Sir David C. Baulcombe at University of Cambridge (*34*). *A. thaliana* lines were obtained from Arabidopsis Biological Resource Center (ABRC) at the Ohio State University. Accession numbers and genetic background were summarized in Table S2.

### Cloning

Nup85 (AT4G32910), Nup88 (AT5G05680), Nup98A (At1g10390), and Nup133 (AT2G05120) were cloned from *Arabidopsis thaliana* genomic DNA. Using the in-fusion cloning strategy, these genomic fragments were inserted into the pEG100 vector with a GFP or a YFP tag. All importin protein constructs (with a Myc-tag) used in this study were reported previously (*12*). To generate the *35S:NPR1-GFP* construct, we purchased CIW00311 from ABRC (Columbus, OH) and used LR clonase (ThermoFisherSci, Waltham, MA) for recombination to generate *35S:NPR1-GFP* using the pBlueScript1510 vector(*71*) (a gift from Dr. Jyan-Chyun Jang at Ohio State University). For adding the SV40 NLS, we used Over-lapping PCR to fuse the NLS to the C-terminal of NPR1 ORF, and the PCR product was inserted into pENTR-D-TOPO vector (ThermoFisherSci) and recombinated into the pBlueScript1510 vector.

PSTVd variants were generated by site-directed mutagenesis using pRZ-Int template and the previously described protocol (*12*). To validate PSTVd progeny sequences, PSTVd was cloned using reverse transcription-PCR followed by cloning into pGEM-T vector (Promega, Madison, WI) as previously described (*12*). Fragment of NPR1 was cloned into pTRV2 vector (ABRC# CD3-1040). pTRV2-NtPDS was used as a negative control (ABRC# CD3-1045).

All clones were verified by Sanger sequencing. Primers are listed in Table S3.

### PSTVd inoculation and RNA gel blots

Plant inoculation and RNA gel blotting procedures were reported previously in details (*13*). Seedlings were inoculated with 150 ng of in vitro transcribed PSTVd RNA, which was evenly applied onto the leaf surface. Several gentle scratches were introduced with pipette tips to facilitate RNA entry into plant tissues. Three weeks post-inoculation, the uppermost fully expanded leaves were collected, and total RNA was extracted using RNAzol (Molecular Research Center, Cincinnati, OH). RNA samples were separated on denaturing urea-PAGE gels and semi-dry transferred onto Hybond membranes (Cytiva, Wilmington, DE), followed by UV crosslinking. Hybridization was performed using specific DIG-labeled riboprobes(*19*), and the membranes were subsequently incubated with alkaline phosphatase–conjugated anti-DIG antibody (Millipore Sigma, Burlington, MA). Chemiluminescence signals were detected using Chemidoc MP (BIO-RAD, Hercules, CA) and quantified using ImageJ (*72*). Statistical analysis was performed using a *t*-test in Prism.

### *Agrobacterium*-mediated *N. benthamiana* transient transformation and immunoblotting

An overnight Agrobacterium culture at 28°C was centrifuged at 6,000 × *g* for 10 min and resuspended in infiltration buffer (10 mM MgCl_₂_, 10 mM MES, pH 5.6, 150 µM acetosyringone) to a final OD_600_ of 0.8. After a 2-h incubation at room temperature, the suspension was infiltrated into fully expanded leaves of *N. benthamiana*. Leaf samples were collected 72 h post-infiltration, ground in liquid nitrogen (∼0.1 g), and extracted with protein extraction buffer (5% SDS, 15% glycerol, 0.3 M DTT, 0.175 M Tris-HCl, pH 8.8, and protease inhibitor cocktail). The homogenate was centrifuged at 12,000 × *g* for 10 min at 4°C, and the supernatant was used as total protein extract. Equal amounts of protein were separated by SDS-PAGE, transferred onto nitrocellulose membranes (ThermoFisherSci), blocked with 5% nonfat milk in TBST, and incubated with primary antibodies followed by HRP-conjugated anti-mouse secondary antibodies (Millipore Sigma).

Nup proteins were detected by monoclonal primary antibody raised in mouse (Santa Cruz Biotechnology Inc.). IMPA proteins were detected using a monoclonal primary antibody against Myc-tag raised in mouse (ThermoFisherSci). ATG3 protein was detected using a polyclonal primary antibody raised in goat (ThermoFisherSci). ATG8 protein was detected using a monoclonal primary antibody raised in rabbit (ThermoFisherSci). Rpb1 was detected using a monoclonal 8W16G antibody raised from mouse (ThermoFisherSci). TFIIIA was detected using a polyclonal antibody raised from rabbit as reported previously (*13*). Protein signals were detected using the Chemidoc MP imaging system and quantified using ImageJ (*72*). *T*-test analysis was performed using Prism.

### Isolation and transformation of *N. benthamiana* protoplasts and fluorescence microscopy

PSTVd infection was first confirmed by RNA gel blots before the plants were used for protoplast assay. The detailed protocol for generating and transfecting protoplasts was published previously (*23*). Fully expanded leaves from 4-6-week-old *N. benthamiana* plants were used for protoplast isolation. The lower epidermis was gently removed with low-residue adhesive tape, and the exposed mesophyll was incubated in enzyme solution (3% Cellulase, 0.8% Macerozyme, 0.5 M mannitol, 20 mM KCl, 10 mM sodium acetate, 20 mM MES, 10 mM CaCl_2_, 5 mM *β*-mercaptoethanol, 0.1% BSA) at 22°C in the dark with gentle agitation for 1 h. The digested suspension was passed through Miracloth, and protoplasts were collected by centrifugation at 100 × *g* for 1 min. The pellet was washed once with W5 solution (154 mM NaCl, 125 mM CaCl_2_, 5 mM KCl, 5 mM MES, pH 5.7), kept on ice for 30 min, and resuspended in MMG solution (0.5 M mannitol, 15 mM MgCl_2_, 4 mM MES) at a density of ∼1-2 × 10^5^ cells/mL. Transformation was carried out using transformation solution (40% w/v PEG4000, 0.2 M mannitol, 0.1M CaCl_2_) for 5 min. After transformation, the reaction was stopped by adding W5 solution. Cells were washed twice by centrifugation, resuspended in an appropriate volume of W5 solution, and incubated in the dark for at least 18 h. For chemical treatments, protoplasts were incubated with 1 mM ASM or 100 µM MG132. Confocal images were acquired using Leica Stellaris 5 confocal microscope equipped with a HC PL APO CS2 40x/1.10 water-immersion objective lens. GFP fluorescence was excited at 488 nm and detected at 497-643 nm. 4’,6-diamidino-2-phenylindole (DAPI) was excited at 405 nm and detected at 420-497 nm. Chlorophyll autofluorescence was excited at 638 nm and detected at 643-839 nm.

### Nuclear and cytoplasmic protein extraction from protoplasts

Protoplasts were collected by centrifugation at 100 × *g* for 1 min and resuspended in extraction buffer I (10 mM Tris-HCl, pH 8.0, 0.4 M sucrose, 0.3% Triton X-100, 5 mM *β*-mercaptoethanol, proteinase inhibitor cocktail). Cells were disrupted by vigorous shaking and incubated on ice for 10 min, followed by centrifugation at 2,100 × *g* for 10 min at 4°C. The supernatant was collected as the cytoplasmic fraction. The pellet was resuspended in extraction buffer II (10 mM Tris-HCl, pH 8.0, 0.25 M sucrose, 1% Triton X-100, 10 mM MgCl_2_, 5 mM *β*-mercaptoethanol, proteinase inhibitor cocktail), incubated on ice for 10 min, and centrifuged at 12,000 × *g* for 10 min at 4°C. After discarding the supernatant, the pellet was washed twice with extraction buffer II and finally resuspended in extraction buffer II to obtain the nuclear fraction. Protein samples were mixed with aforementioned protein extraction buffer, boiled for 5 min, and subjected to immunoblotting after a brief spin.

### Chemical treatments

At one week post PSTVd infection, plants were subjected to a seven-day foliar spraying treatment. The chemicals, including ASM (Santa Cruz Biotechnology Inc., Dallas, TX), echinocystic acid (Millipore Sigma), and sterile water as control, were evenly applied to the leaf surface. For ASM, 100 µM solution was used based on a previous report(*73*). For echinocystic acid, we used recommended 4.23 µM concentration for spraying(*61*). Beginning in the third week after infection, the uppermost fully expanded leaves were collected weekly for RNA extraction or subsequent analyses. MG132 (Cayman Chemical, Ann Arbor, MI) was infiltrated into leaves with a concentration of 50 µM 4 hr before sample collection.

### Proteomics analysis

Label-free proteomics analysis was performed using a fee-based service provided by Creative Proteomics (Shirley, NY). According to their provided protocol, leaf samples were homogenized in lysis buffer (100 mM Tris-HCl, pH 8.0, 8 M urea, 1% protease inhibitor) and centrifuged at 12,000 x *g* for 10 min at 4°C. Protein concentrations were determined by BCA assay, and 120 µg of protein per sample was reduced with 10 mM TCEP at 56 °C for 1 h, alkylated with 20 mM IAA in the dark for 30 min, and precipitated with six volumes of cold acetone at –20°C overnight. The protein pellets were resuspended in 100 mM TEAB and digested with trypsin (1:50, w/w) at 37°C overnight. Peptides were dried and reconstituted in 0.1% formic acid prior to LC-MS/MS analysis. Peptides (1 µg) were loaded onto a trapping column (PepMap C18, 100 µm × 2 cm) and separated on an analytical column (PepMap C18, 75 µm × 50 cm) using an Ultimate 3000 nano UHPLC system (ThermoFisherSci) with a 90-min gradient (2-40% acetonitrile, 0.1% formic acid) at 250 nL/min. Mass spectra were acquired on an Orbitrap mass spectrometer in data-dependent mode (full scan, m/z 300-1650, resolution 60000; Top20 MS/MS, resolution 15000; NCE 28%; dynamic exclusion 30 s). Raw MS files were searched against the ITAG4.0 protein database using MaxQuant (v1.6.2.6) with default parameters: trypsin specificity, up to two missed cleavages, carbamidomethylation (C, fixed), oxidation (M, variable), precursor mass tolerance 10 ppm, and fragment mass tolerance 0.02 Da.

### Salicylic Acid content analysis

Tomato plants were confirmed to be infected with PSTVd before sampling. Approximately 200 mg of tissue was collected from each biological replicate and ground to a fine powder in liquid nitrogen. The ground tissue was extracted with 600 µL of 90% (v/v) methanol in water. Samples were sonicated on ice for 15 minutes and centrifuged at 15,000 × *g* for 5 minutes. 450 µL of the supernatant was then collected, and the remaining sample material was re-extracted with 600 µL of 100% methanol. Following a second round of sonication and centrifugation, 900 µL of supernatant was collected. The combined extract was supplemented with 50 µL of 0.2 M sodium hydroxide and dried to completion using a SpeedVac. The dried residue was resuspended in 500 µL of 5% (v/v) trichloroacetic acid (TCA) in water, sonicated for 15 minutes on ice, and centrifuged at 15,000 × *g* for 5 minutes. 450 µL of the supernatant was collected and extracted with 500 µL of ethyl acetate-cyclopentane (1:1), followed by sonication on ice for 15 minutes and centrifugation at 15,000 × *g* for 5 minutes. 420 µL of the organic phase was collected. The ethyl acetate extraction was repeated, and a final volume of 920 µL was collected. Subsequently, 60 µL of 10% (v/v) methanol in 0.2 M acetate buffer (pH 5.5) was added, and the samples were dried. The dried extracts were resuspended in 150 µL of 99% methanol (v/v) in water for HPLC analysis. For analysis of SAG, the residual sample material was hydrolyzed with 12N HCl at 80°C for 60 minutes prior to SA extraction as described above. The samples were then analyzed using a Vanquish HPLC system equipped with an Acclaim^TM^ RSLC 120 C18 column (2.2 µm, 120Å, 3×100 mm). Solvent A consisted of 10% methanol in 0.2 M acetate buffer (pH 5.5) (v/v) and solvent B consisted of 99% methanol in water (v/v). The gradient program was as follows: 5% B at 0 min, 80% B at 8.5 min, 95% B at 9 min, and 95% B at 11.5 min. The flow rate was set to 0.7 mL/min. Fluorescence detection was performed at an excitation wavelength of 295 nm and an emission wavelength of 407 nm. Peak areas were integrated and SA concentrations were determined using a salicylic acid standard curve. SA levels were normalized to the fresh weights of the tissues used for extraction.

### *In vivo* smStructure-seq library construction

Dimethyl sulphate (DMS) was used to generate *in vivo* smStructure-seq libraries as previously described (*74*). Briefly, harvested leaves were submerged in 20 mL of 1× DMS reaction buffer (200 mM potassium acetate, 200 mM Bicine (pH 8.0), and 5 mM MgCl_₂_) in 50 mL Falcon tubes. Leaves were incubated with 2% DMS for 15 min and with shaking at 800 rpm. The reaction was quenched with freshly prepared dithiothreitol, after which samples were washed with deionized water, immediately frozen in liquid nitrogen, and ground into powder. Total RNA was extracted using the RNeasy kit (Qiagen, Manchester, UK), followed by DNase I treatment according to the manufacturer’s instructions. Control samples (minus) were prepared using water instead of DMS, following the same procedure. Reverse transcription was performed using Induro Reverse Transcriptase (NEB) with 1 *μ*L of RNase inhibitor. The reaction was incubated at 60°C for 180 min. PCR amplification was performed for 20 cycles using two pairs of specific primers (Table S3) with Q5 High-Fidelity DNA Polymerase (NEB). Two independent biological replicates were generated for the plus libraries. Purified DNA samples were subsequently used for Nanopore library preparation following the manufacturer’s instructions for the native barcoding kit (NBD-24).

### smStructure-seq data processing and structural analyses

Raw reads from both DMS-treated (plus) and untreated (minus) samples were basecalled and demultiplexed using Dorado (https://github.com/nanoporetech/dorado), and were aligned to the PSTVd transcriptome using minimap2 (*75*) with the parameters -t 8 --secondary=no -G 12000. DMS reactivity was calculated from mutation rates using the RNAFramework pipeline (*76*). Mutation rates at each nucleotide position were derived as the ratio of mutation counts to total read coverage at each nucleotide position, followed by background correction using the corresponding minus control. The resulting raw reactivities were normalized using the 2–8% normalization method to ensure comparability across samples (*76*).

The Gini index was calculated based on normalized DMS reactivity values to assess the unevenness of reactivity distribution across the RNA, following the implementation in the RNAFramework pipeline (*76*). RNA secondary structure models were generated using RNAfold (ViennaRNA package) (*77*), incorporating normalized DMS reactivities as soft constraints.

### DiffScan analysis

Differential RNA structure analysis between conditions was performed using DiffScan (*39*), taking normalized DMS reactivity values as input. DiffScan evaluates position-level differences between conditions using a nonparametric statistical test within local nucleotide windows (Wilcoxon test) and subsequently aggregates these signals across regions using scan statistics to identify structurally variable regions (SVRs) with statistical significance.

### Structural analysis by DaVinci

The DaVinci pipeline was performed largely as previously described, with minor modifications (*74*). Briefly, per-read mutation profiles were converted into individual bit vectors and used for constraint-guided RNA folding, generating one RNA secondary structure per molecule. To reduce background noise, positions with mutation rates in the untreated (minus) sample equal to or greater than those in the DMS-treated (plus) sample were excluded from constraint incorporation, and G-to-A substitutions were specifically filtered out during bit-vector generation. RNA structures were represented in dot-bracket notation and further transformed into RNA structural elements using the rnaConvert function from the Forgi package. Shannon entropy and R5 region similarity analysis were performed at the single-molecule level based on the resulting RNA structures.

### Structural similarity analysis of the R5 region

To quantify structural similarity of the R5 region under different conditions, each single-molecule RNA structure was compared with the reference R5 structure based on base-pairing patterns. For each molecule, base pairs in the observed structure were extracted from the dot-bracket notation and compared with those in the reference structure. Three complementary metrics were used: F1 score, Jaccard index, and helix integrity score.

### F1 score

The F1 score (*78*) was used to evaluate overall agreement between the observed structure and the reference structure at the base-pair level. Base pairs were classified as follows: true positives (TP), base pairs present in both the observed and reference structures; false positives (FP), base pairs present only in the observed structure; and false negatives (FN), base pairs present only in the reference structure. The F1 score was calculated as:

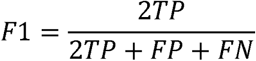

This metric ranges from 0 to 1, with higher values indicating greater similarity to the reference R5 structure.

### Jaccard index

The Jaccard index (*79*) was used to measure the overlap between the observed and reference base-pair sets. It was calculated as:

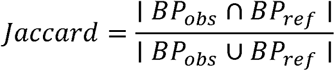

where *BP_obs_* and *BP_ref_* represent the base-pair sets of the observed from each single-molecule RNA structure and reference structures, respectively. This metric also ranges from 0 to 1, with higher values indicating a greater proportion of shared base pairs.

### Helix integrity score

The helix integrity score was used to assess structural conservation at the helix level. This metric is based on helix-wise evaluation frameworks commonly used in RNA secondary structure benchmarking (*80*). In the reference structure, base pairs were grouped into helices, defined as consecutive nested base pairs (e.g., (*i, j*), (*i+ 1, j - 1*)), representing continuous stems.

For each reference helix, the fraction of its base pairs retained in the observed structure was calculated. A helix was considered intact if at least 50% of its base pairs were present in the observed structure.

The helix integrity score was then defined as:

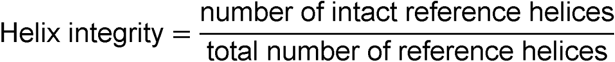

This metric is equivalent to helix-level sensitivity (i.e., the fraction of correctly recovered helices) and ranges from 0 to 1, reflecting preservation of the overall helical scaffold of the reference R5 structure.

### Statistical analysis

Differences between conditions were evaluated at the single-molecule level. Distributions of F1 score, Jaccard index, and helix integrity score were compared using the two-sided Mann–Whitney U test, and exact *P* values were reported.

### Data, Materials, and Software Availability

Previously published RNA-Seq data (NCBI PRJNA353731) were used for this work(*27*). Proteomics data and nanopore sequencing data have been summarized in figures and tables, and raw data will be publicly available upon acceptance. All other data are included in the manuscript and/or supporting information.

## ACKNOWLEDGMENTS

We apologize to colleagues whose work was not cited due to the page limits. We are in debt to Drs. Yiling Fang and Yangnan Gu at University of University of California, Berkeley, for sharing their expertise on nuclear pore complex and constructs of Nup proteins. This work is supported by US National Science Foundation (MCB-2350392 and IOS-2410009) to Ying Wang, and (IOS-2142898) to Jeongim Kim, and UK Biotechnology and Biological Sciences Research Council (BB/Y002849/1) to Yiliang Ding. Yunhan Wang is partially supported by CALS Dean’s scholarship from University of Florida.

## Figure legends

**Figure S1.** Reproducibility of mutation rates across biological replicates at different temperatures.

**Figure S2.** Summary of published multi-probing analysis of loop 24.

**Figure S3.** RNA gel blot analysis confirmed PSTVd infection in samples used for SA content measurement.

**Figure S4.** RNA gel blot confirmed PSTVd Infection for proteomics analysis. **Figure S5.** Subcellular localization of NPR1 and NPR1-NLS in infected protoplasts. **Figure S6.** Multiple probing data revised the model of PSTVd loop24.

**Table S1.** The 201 position nucleotides in PSTVd variants.

**Table S2.** Plant materials used in this study.

**Table S3.** Primer sequences.

**Dataset S1.** Proteomics data summary.

**Dataset S2**. Comparison of RNA-Seq and proteomics data of nucleoporins and importins.

