## Supplementary figures and images for "Elevated temperature enhances the infectivity of a pathogenic noncoding RNA and the likelihood of its host expansion"

### Fig.S1

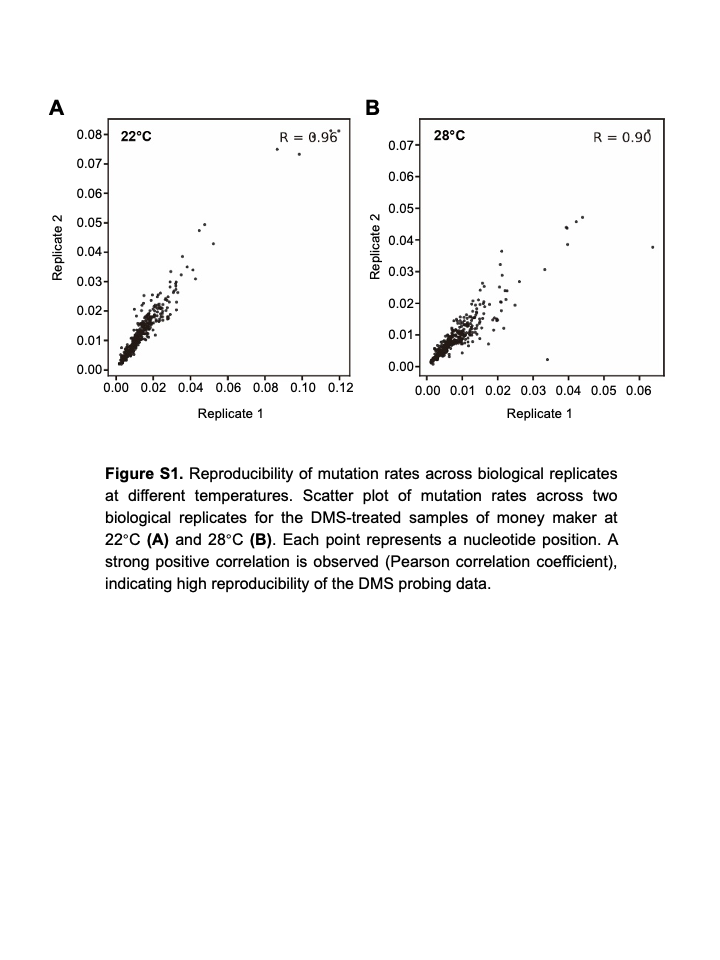

### Fig.S2

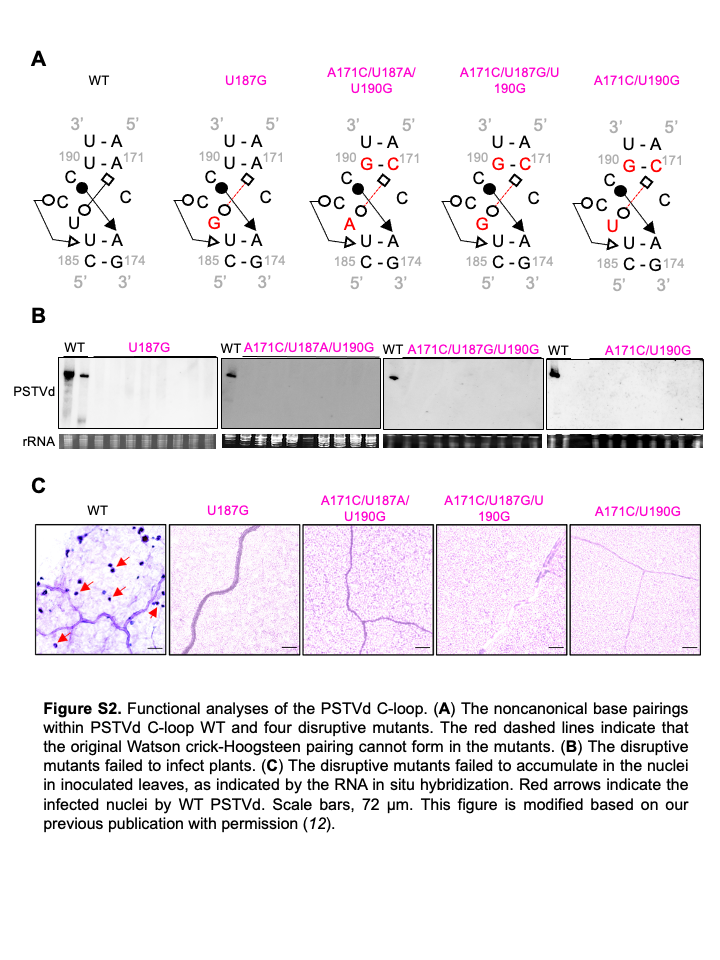

### Fig.S3

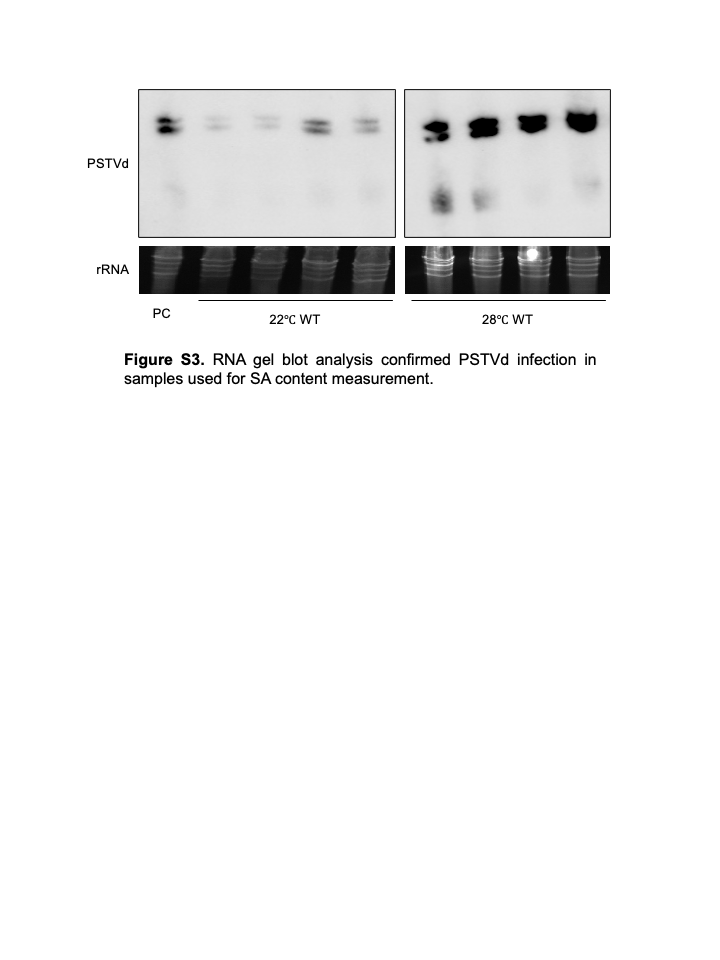

### Fig.S4

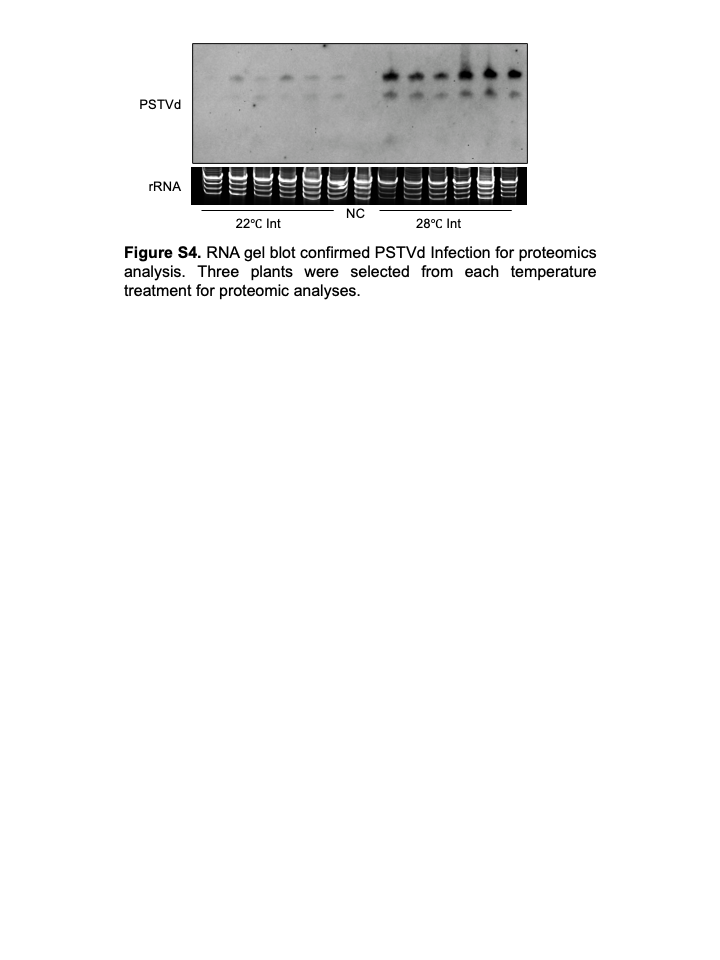

### Fig.S5

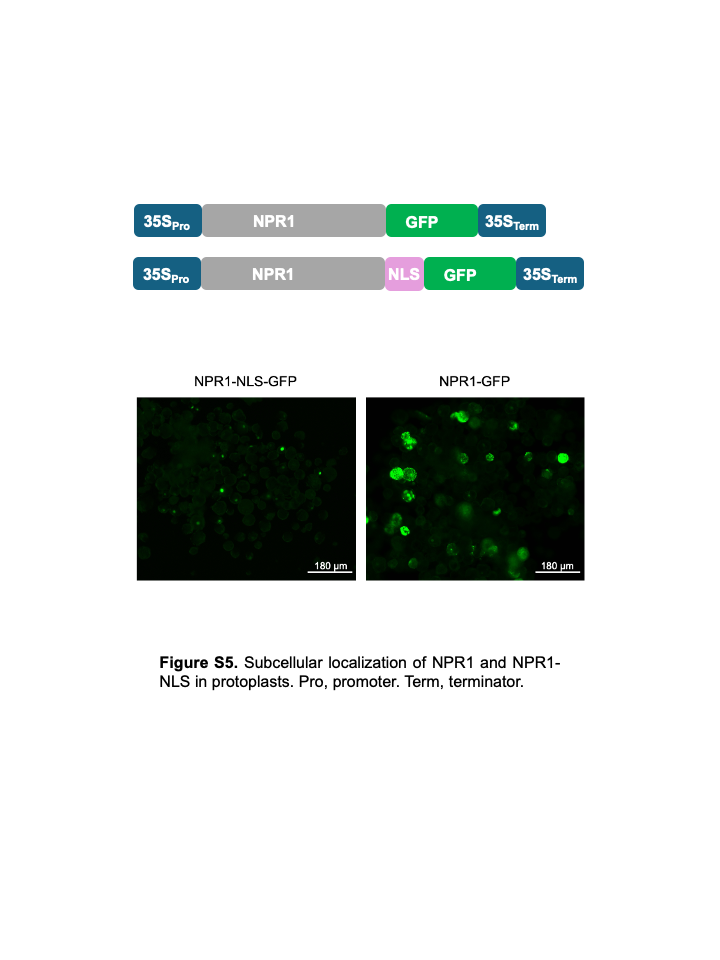

### Fig.S6

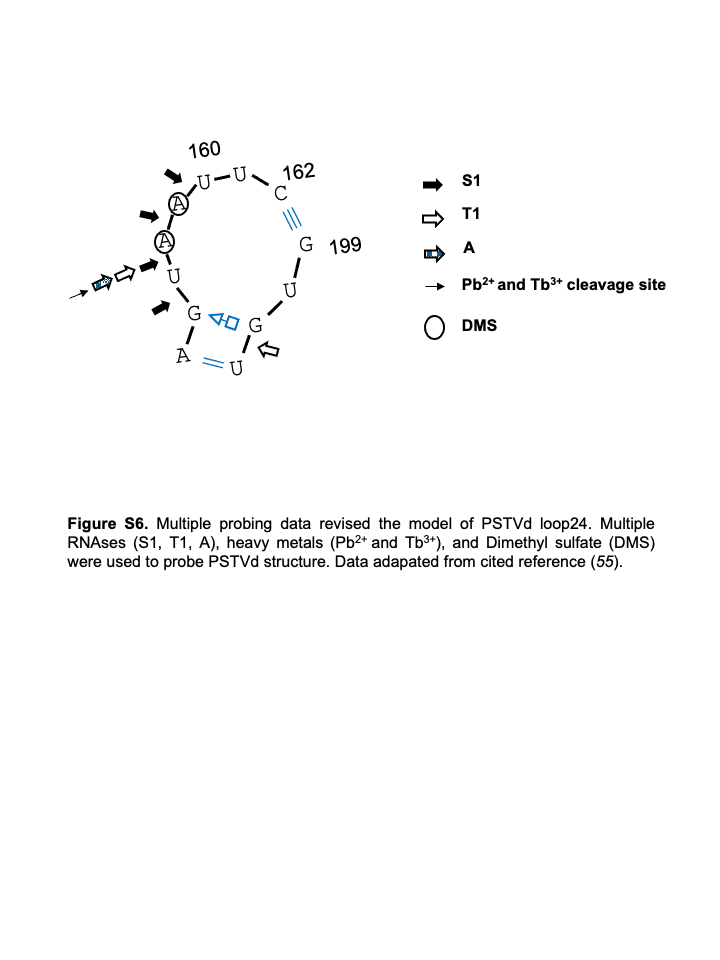
